## Supplementary Figures for "Context-dependent activation and evolutionary buffering of a mating pheromone in fission yeast"

Supplementary Fig. S1

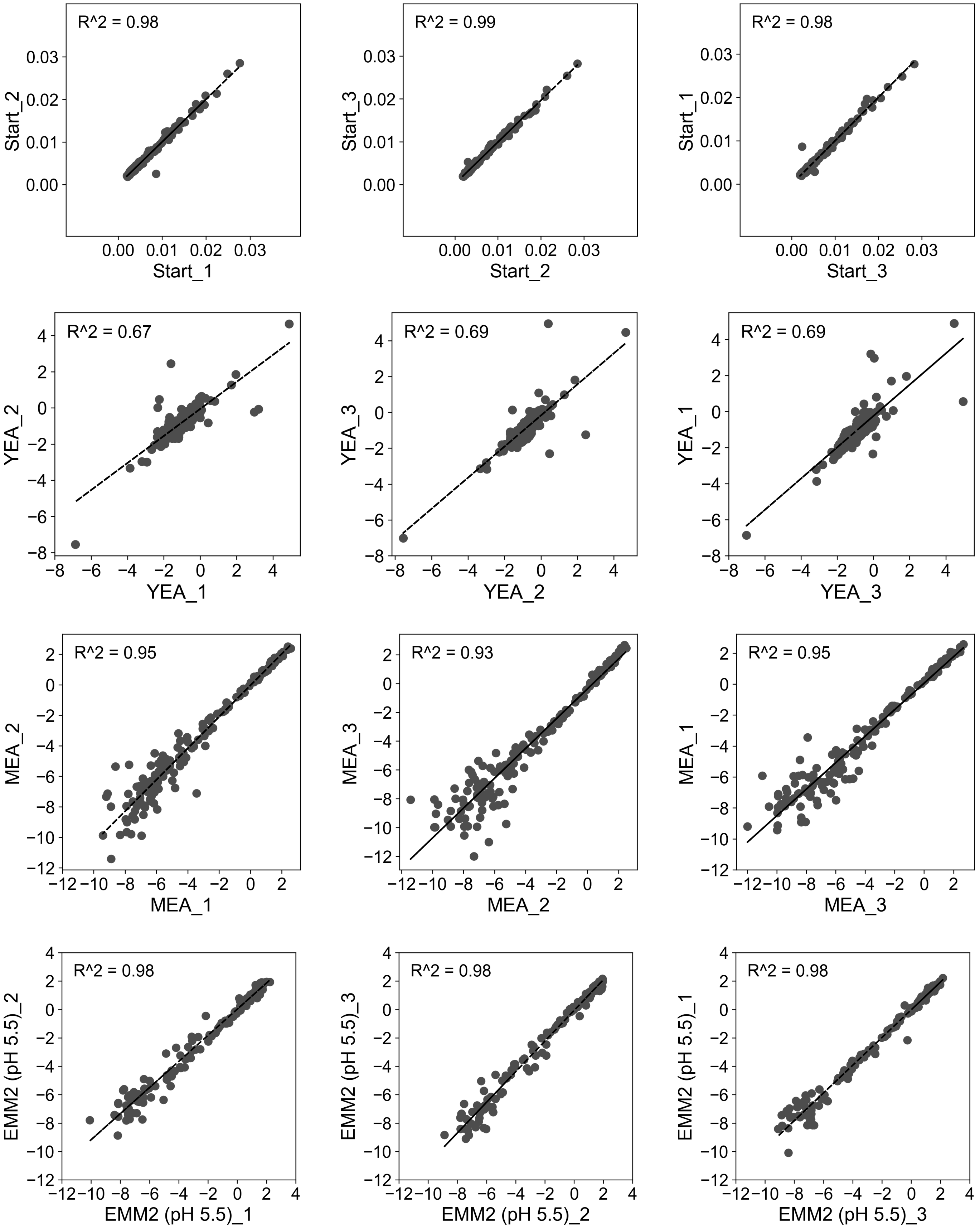

Supplementary Fig. S1 (continued)

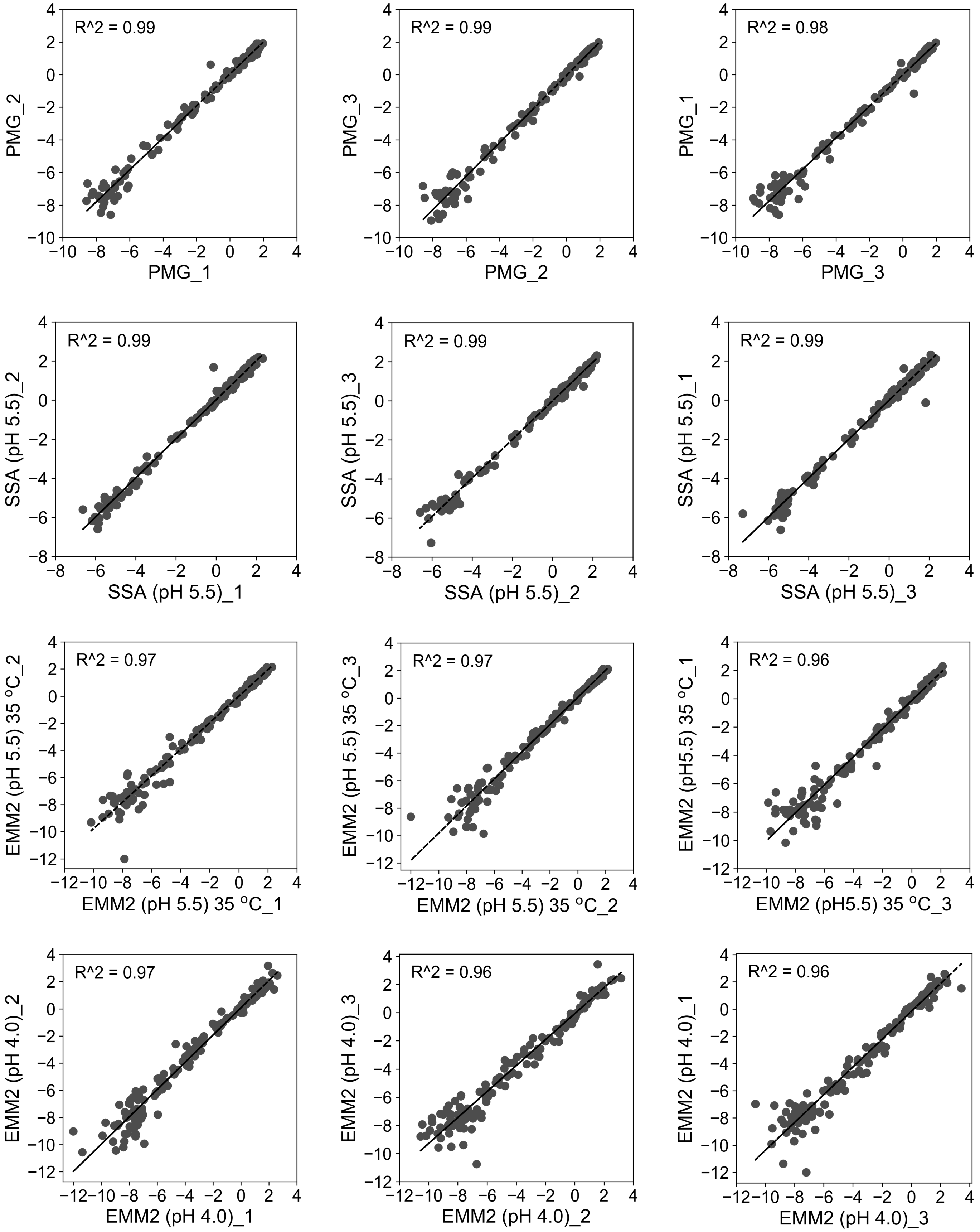

Supplementary Fig. S1 (continued)

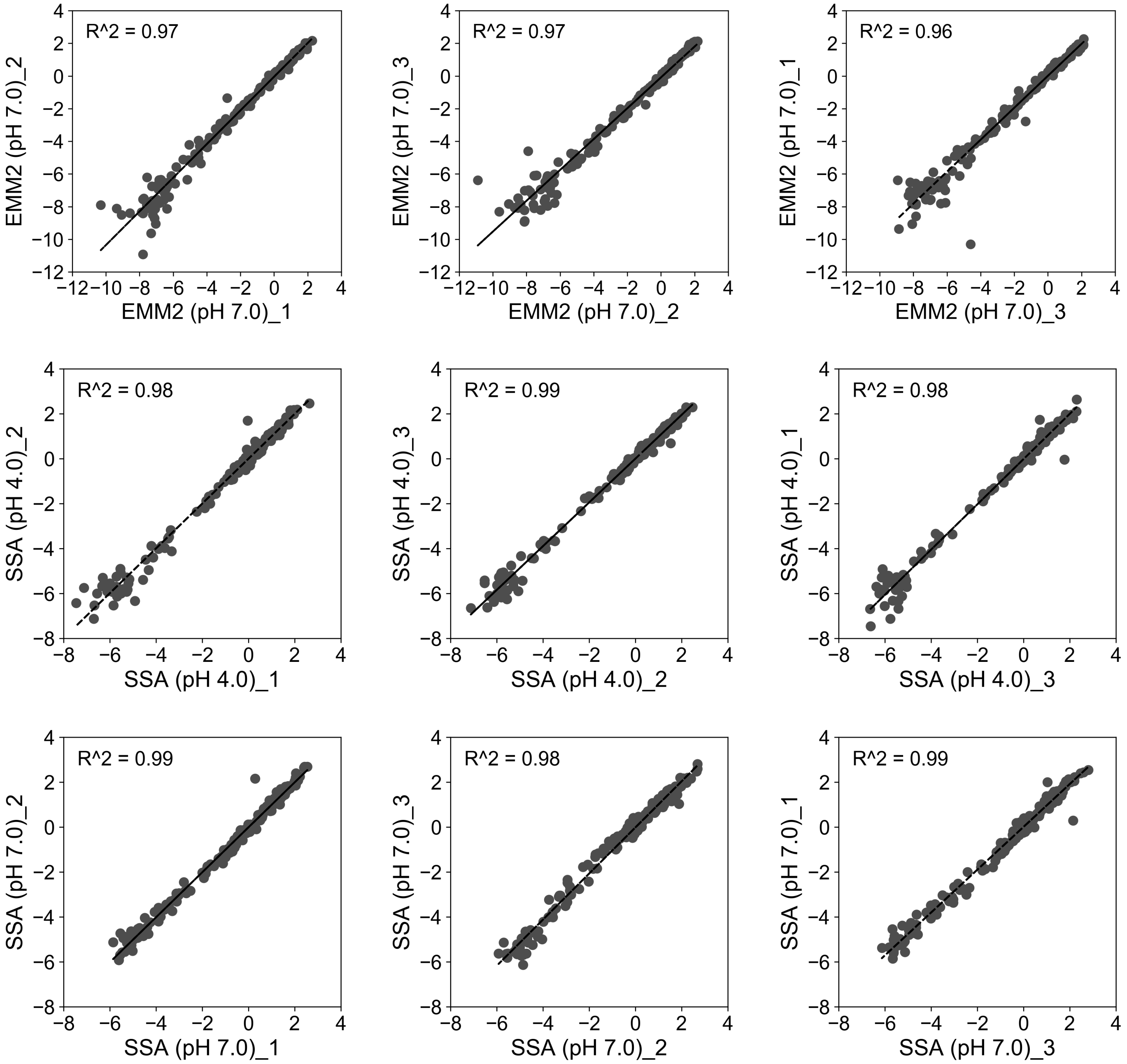

Supplementary Fig. S2

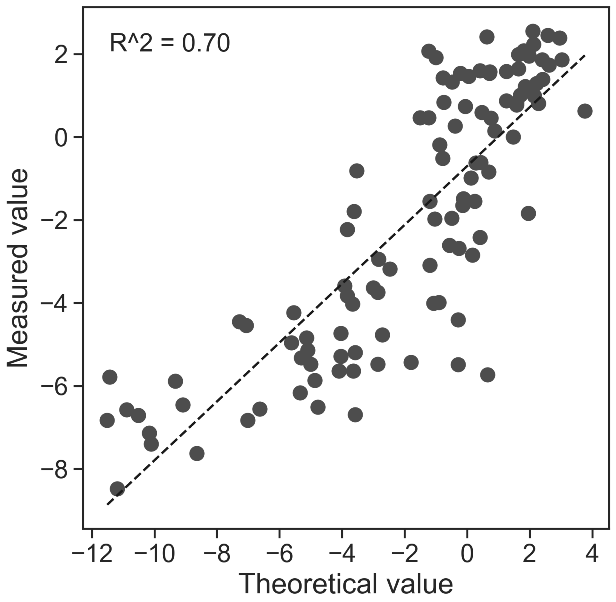
